# The Ptk2-Pma1 pathway enhances tolerance to terbinafine in *Trichophyton rubrum*

**DOI:** 10.1101/2023.12.07.570643

**Authors:** Masaki Ishii, Tsuyoshi Yamada, Michel Monod, Shinya Ohata

**Affiliations:** Research Institute of Pharmaceutical Sciences, Faculty of Pharmacy, Musashino University, Tokyo, Japan; Teikyo University Institute of Medical Mycology, Teikyo University, Tokyo, Japan; Asia International Institute of Infectious Disease Control, Teikyo University, Tokyo, Japan; Department of Dermatology, Centre Hospitalier Universitaire Vaudois, Lausanne, Switzerland; Faculty of Biology and Medicine, University of Lausanne, 1015 Lausanne, Switzerland

**Author notes:** Address correspondence to: M. Ishii, S. Ohata. **Competing interests:** The authors declare no competing interests.

**Keywords:** Dermatophytosis, terbinafine resistance, proton pump, Pma1, Ptk2, omeprazole, *Trichophyton rubrum*

## Abstract

The increasing prevalence of dermatophyte resistance to terbinafine, a key drug in the treatment of dermatophytosis, represents a significant obstacle to treatment. *Trichophyton rubrum* is the most commonly isolated fungus in dermatophytosis. In *T. rubrum*, we identified TERG_07844, a gene encoding a previously uncharacterized putative protein kinase, as an ortholog of budding yeast *Saccharomyces cerevisiae* polyamine transport kinase 2 (Ptk2) and found that *T. rubrum* Ptk2 (TrPtk2) is involved in terbinafine tolerance. In both *T. rubrum* and *S. cerevisiae*, Ptk2 knockout strains were more sensitive to terbinafine compared to the wild types, suggesting that promotion of terbinafine tolerance is a conserved function of fungal Ptk2. The *T. rubrum* Ptk2 knockout strain (ΔTrPtk2) was sensitive to omeprazole, an inhibitor of plasma membrane proton pump Pma1, which is activated through phosphorylation by Ptk2 in *S. cerevisiae*. Overexpression of *T. rubrum* Pma1 (TrPma1) in ΔTrPtk2 suppressed terbinafine sensitivity, suggesting that the induction of terbinafine tolerance by TrPtk2 is mediated by TrPma1. Furthermore, omeprazole increased the terbinafine sensitivity of clinically isolated terbinafine-resistant strains. These findings suggest that, in dermatophytes, the TrPtk2-TrPma1 pathway plays a key role in promoting intrinsic terbinafine tolerance and may serve as a potential target for combinational antifungal therapy against terbinafine-resistant dermatophytes.

## Introduction

Dermatophytes are fungal pathogens that infect the surface tissues of mammals and other animals, causing symptoms such as itching and nail deformities. Dermatophytes can also exacerbate allergies in patients with asthma, significantly reducing their quality of life (1, 2). Antifungal drugs have been developed for fungal infections, and terbinafine, with its ability to inhibit squalene epoxidase in the ergosterol synthesis pathway, has been a highly effective medicine. However, terbinafine-resistant fungi have emerged in recent years (3–5), and the prevalence of resistant strains is a serious concern for the future treatment of dermatophytosis.

Recently, we reported that resistance to terbinafine is caused by mutations in squalene epoxidase, the target of terbinafine (6). The L393F and F397L mutations in squalene epoxidase are the major causes of terbinafine resistance reported worldwide (3, 6–8). However, therapeutic targets and compounds to alleviate this resistance remain to be identified.

Drug repurposing is the practice of using a therapeutic agent that is already approved for the treatment of another disease. In the field of infectious diseases, drug repurposing has become a significant research strategy to discover effective therapeutics against drug-resistant bacteria (9, 10). Previous reports encourage the application of drug repurposing to aid in the research and development of therapies for dermatophytosis.

In the present study, we found that ablation of the gene encoding the putative protein kinase TERG_07844 in *Trichophyton rubrum* resulted in the decrease of tolerance to terbinafine. Our findings suggest that TERG_07844 gene product has a homologous function to the protein kinase Ptk2 of budding yeast and that the proton pump Pma1 functions downstream of TERG_07844 gene product in terbinafine tolerance. We also found that omeprazole, a proton pump inhibitor approved for clinical use, potentiated the antifungal effect of terbinafine in terbinafine-resistant isolates. These results suggest that the Ptk2-Pma1 pathway enhances resistance to terbinafine in *Trichophyton rubrum* and could be a potential target for antifungal treatment.

## Results

### TERG_07844 is involved in terbinafine tolerance

Protein kinases are involved in a wide range of physiological activities, including the regulation of intracellular ion concentrations and responses to external stresses such as antifungal drugs (11). Whole-genome analyses of dermatophytes have revealed a large number of genes encoding kinases of unknown function (12). Among these genes, we focused on TERG_07844 because it is conserved among dermatophytes and is highly expressed in *T. rubrum* (13–15). We generated a TERG_07844 knockout stain from the terbinafine- susceptible *T. rubrum* strain CBS118892 by replacing the TERG_07844 open reading frame (ORF) with the neomycin resistance gene (*nptII*) cassette (ΔTERG_07844, Figure 1A). We also generated a revertant strain (eYFP-TERG_07844C) by random integration of the *eYFP*- TERG_07844 gene, which expresses TERG_07844 gene product (XP_047604827) tagged with enhanced yellow fluorescent protein (eYFP) at the N-terminus (eYFP- XP_047604827), in the genome of ΔTERG_07844.

**Figure 1.**
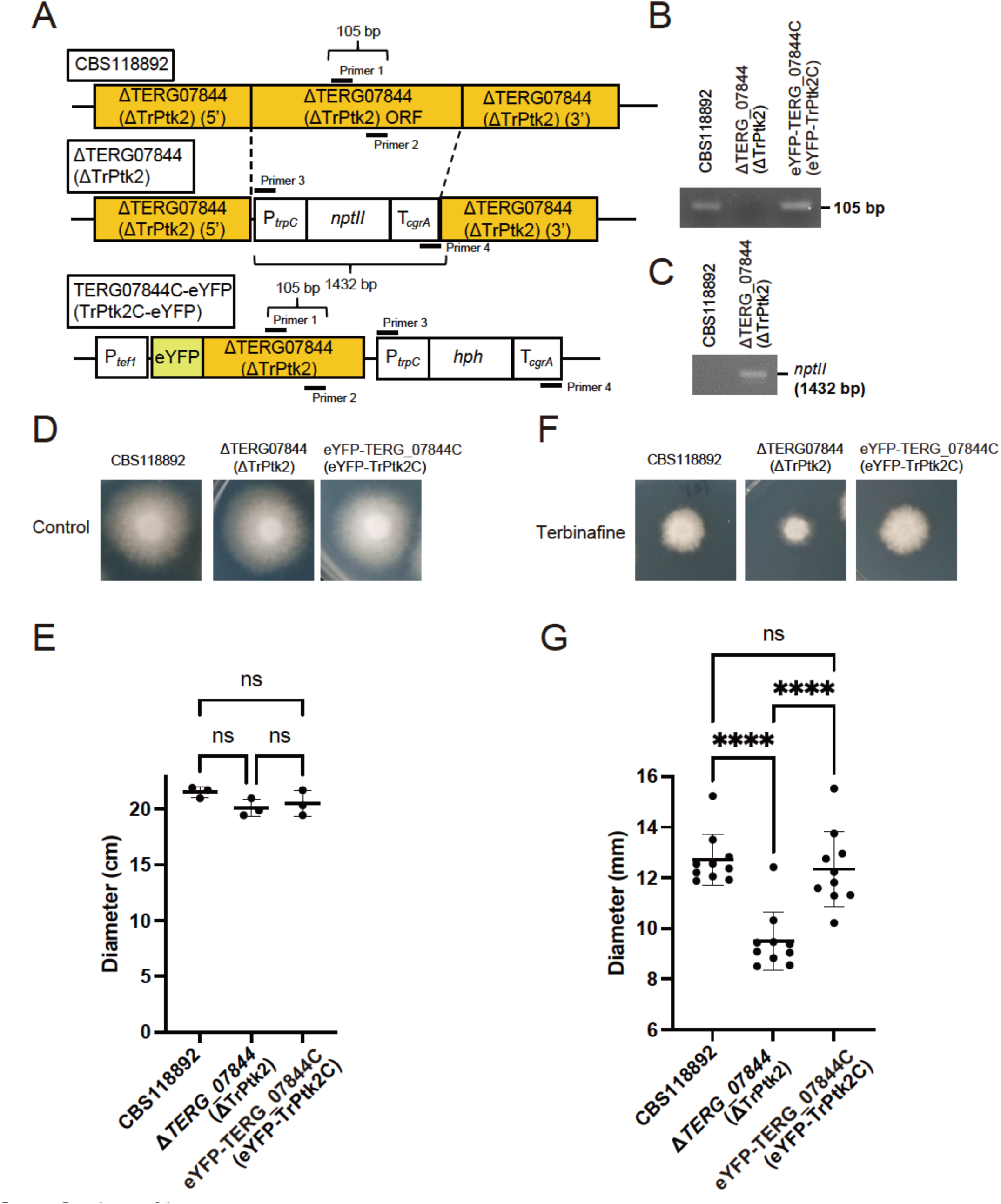
Contribution of the TERG_07844 (TrPtk2) gene to terbinafine sensitivity in *Trichophyton rubrum*. (A) Schematic representation of the TERG_07844 (TrPtk2) wild-type allele (top, CBS118892), deletion construct (middle, ΔTERG_07844), and revertant construct (bottom, eYFP-TERG_07844C). (B) PCR analysis of CBS118892 and ΔTERG_07844 (TrPtk2) and eYFP-TERG_07844C (eYFP-TrPtk2C) using primer pairs 1 and 2. (C) PCR analysis of CBS118892 and ΔTERG_07844 (TrPtk2) and eYFP-TERG_07844C (eYFP-TrPtk2C) using primer pairs 3 and 4. (D–G) Terbinafine susceptibility of CBS118892, ΔTERG_07844, and eYFP-TERG_07844C, in the presence and absence of low concentrations of terbinafine. (D) Spores of CBS118892, ΔTERG_07844 (TrPtk2), and eYFP-TERG_07844C (eYFP-TrPtk2C) were inoculated on RPMI 1640 for 14 days. (E) The diameter of the mycelium on RPMI 1640 after 14 days was measured. The dots on the graph represent the diameter of individual samples (n = 3). (F) Spores of CBS118892, ΔTERG_07844 (TrPtk2), and eYFP-TERG_07844C (eYFP-TrPtk2C) were inoculated on RPMI 1640 with 5 ng/mL of terbinafine for 14 days. (G) The diameter of the mycelium on RPMI 1640 with 5 ng/mL terbinafine after 14 days was measured. The dots on the graph represent the diameter of individual samples (n = 10).

To confirm the loss of the TERG_07844 ORF, PCR was performed using primer pairs designed within the TERG_07844 ORF (primers 1 and 2 in Figure 1A) and the neomycin resistance gene *nptII* cassette (primers 3 and 4 in Figure 1A). PCR using the former primer pair amplified the PCR products in the parental strain CBS118892 and eYFP-TERG_07844C, but not in ΔTERG_07844 (Figure 1B). Conversely, PCR using the latter primer pair did not amplify the PCR products in CBS118892 but did in ΔTERG_07844 (Figure 1C).

To analyze the terbinafine susceptibility of *T. rubrum* CBS118892, ΔTERG_07844, and eYFP-TERG_07844C, we cultured these strains on agar plates in the presence and absence of low concentrations of terbinafine (Figure 1D and F) and measured the diameter of the colonies (Figure 1E and G). The mycelial growth of ΔTERG_07844 was comparable to that of CBS118892 and eYFP-TERG_07844C on the agar medium without terbinafine (Figure 1D and E). However, on agar medium containing terbinafine, the mycelial growth of ΔTERG_07844 was significantly reduced (Figure 1F and G). These results suggest that TERG_07844 is involved in terbinafine tolerance in *T. rubrum*.

### XP_047604827 encoded by TERG_07844 in *T. rubrum* is phylogenetically and functionally similar to *S. cerevisiae* Ptk2

To gain insight into TERG_07844, we performed a phylogenetic tree analysis to determine which kinases in *S*. *cerevisiae* are similar to XP_047604827 encoded by TERG_07844 (Figure 2A). The phylogenetic tree revealed that XP_047604827 is grouped with the halotolerance kinases Sat4 (accession number NP_009934) and Hal5 (accession number NP_012370) from *S*. *cerevisiae*. Deficiencies in these kinases result in the decrease of high salt tolerance in *S*. *cerevisiae* (16). The polyamine transport kinase, Ptk2 (accession number NP_012593), was also found in proximity to XP_047604827. In contrast to Sat4 and Hal5, the absence of Ptk2 has been reported to cause high salt tolerance in *S*. *cerevisiae* (17, 18). Near XP_047604827, the protein XP_964224, identified as a Ptk2 ortholog in the filamentous fungus *Neurospora crassa* (19), was also found (Figure 2A). To determine whether XP_047604827 is functionally related to either Sat4/Hal5 or Ptk2, we examined the response of *T. rubrum* ΔTERG_07844 in a medium containing high salt concentrations. Compared to the terbinafine-sensitive strain CBS118892, ΔTERG_07844 exhibited enhanced mycelial growth in the presence of 0.5 M NaCl and displayed high salt tolerance, like the Ptk2-deficient *S. cerevisiae* strain ΔScPtk2 (17, 18). Moreover, the sensitivity of ΔTERG_07844 to compounds to which ΔScPtk2 is resistant was investigated (18). The results showed that ΔTERG_07844 is resistant to spermine, lithium chloride, and hygromycin (Figure 2B and 2C). These salt tolerances were significantly reduced in eYFP-TERG_07844C (Figure 2B and 2C). These results suggest that XP_047604827 encoded by TERG_07844 has phylogenetic and functional similarities to the Ptk2 protein of budding yeast. Consequently, we refer to *T. rubrum* XP_047604827 encoded by TERG_07844 as TrPtk2 in this study.

**Figure 2.**
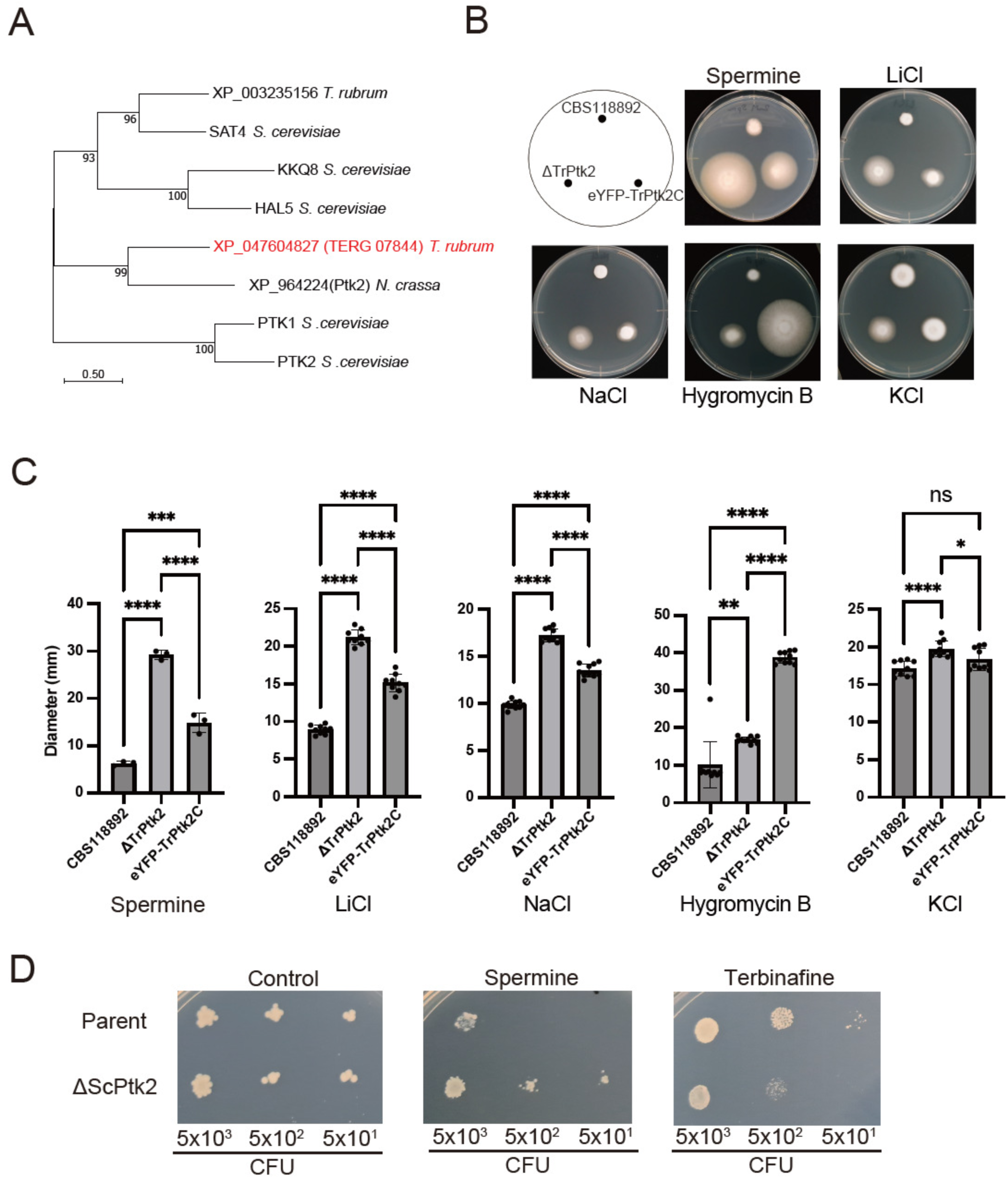
TERG_07844 gene product XP_047604827 of *T. rubrum* is phylogenetically and functionally similar to Ptk2. (A) Phylogenetic tree of fungal proteins related to XP_047604827 (TrPtk2) encoded by TERG_07844 inferred using the maximum likelihood method. The optimal tree is displayed. Evolutionary distances were estimated using the JTT model. (B–C) Effect of spermine, various salts, and hygromycin B on TERG_07844 (TrPtk2) growth. Spores of CBS118892, ΔTERG_07844 (TrPtk2), and eYFP-TERG_07844C (eYFP-TrPtk2C) were inoculated on SDA with 2 mM spermine, 50 mM LiCl, 0.5 M NaCl, 100 μg/mL hygromycin B, and 0.5 M KCl and incubated for 14 days (B). The diameter of the mycelium was measured (C). The dots on the graph represent the diameter of individual samples (n = 3 for spermine; n = 10 for others). The hygromycin resistance gene (*hph*) was used as a selectable marker to isolate eYFP-TERG_07844C (eYFP-TrPtk2C), and this strain was found to be resistant to hygromycin (B and C). (D) Acquired resistance of *S. cerevisiae* to spermine and terbinafine after deletion of the gene encoding Ptk2. Parent and ΔScPtk2 were grown in synthetic defined medium, and serial dilutions were dropped on synthetic defined agar plates with 2 mM spermine or 50 μg/mL terbinafine. Growth was measured after 3 days.

To investigate the general impact of fungal Ptk2 on terbinafine resistance, we assessed the sensitivity of the Ptk2-deficient *S. cerevisiae* strain ΔScPtk2 (Table 1) to terbinafine. As previously reported,ΔScPtk2 was resistant to spermine (Figure 2D left and center panels). Interestingly, ΔScPtk2 was sensitive to terbinafine (Figure 2D right panel), similar to ΔTrPtk2 (Figure 1G). These observations suggest that the contribution of fungal Ptk2 to terbinafine tolerance is evolutionarily conserved in fungi.

**Table 1.** Fungal strains used in this study.

| Species and strains | Description | Reference |
| --- | --- | --- |
| *Trichophyton rubrum* |  |  |
| **CBS118892** | A Terbinafine-sensitive clinical isolate from a patient nail sample. | (12) |
| **ΔTERG_07844 (ΔTrPtk2)** | The TERG_07844 open reading frame (ORF) was replaced with the neomycin resistance gene (*nptII*) in the strain. This strain was derived from CBS118892. | This study |
| **eYFP-TERG_07844C (eYFP -TrPtk2C)** | A complementary (revertant) strain by random integration of the N-terminal enhanced yellow fluorescent protein (eYFP) tag-fused TERG_07844 gene into the ΔTERG_07844 (ΔTrPtk2) genome. | This study |
| **TrPma1OE-eYFP** | ΔTrPtk2 overexpressing eYFP-tagged TrPma1 by random integration of the C-terminal enhanced yellow fluorescent protein (eYFP) tag-fused TrPma1 gene into the ΔTERG_07844 (ΔTrPtk2) genome. | This study |
| **TIMM20092** | A terbinafine-resistant clinical isolate has a F397L in the squalene epoxidase. | (6) |
| **TIMM20093** | A terbinafine-resistant clinical isolate has a L393F in the squalene epoxidase. | (6) |
| **TIMM20094** | A terbinafine-resistant clinical isolate has a L393F in the squalene epoxidase. | (6) |
| *Saccharomyces cerevisiae* |  |  |
| **Parent** | A strain harboring pYES2-HTH derived from BY4741 (purchased from Horizon Discovery Ltd.). | This study |
| **ΔScPtk2** | A strain harboring pYES2-HTH derived from YJR059W(a Ptk2 deletion strain derived from BY4741) (purchased from Horizon Discovery Ltd.). | This study |

### Overexpression of TrPma1 suppresses the terbinafine sensitivity of ΔTrPtk2

In *S. cerevisiae*, the proton pump Pma1 is an essential protein for fungal growth (20). Pma1 is the most established substrate of Ptk2 and is activated by this kinase through phosphorylation *in S. cerevisiae* (21, 22). To investigate whether *T. rubrum* Pma1 (TrPma1) functions downstream of TrPtk2 in *T. rubrum*, we assessed the sensitivity of the strain ΔTrPtk2 to omeprazole, an inhibitor of Pma1 in the yeast (23). On omeprazole-free agar medium, growth of CBS118892 (parent), ΔTrPtk2, and eYFP-TrPtk2C (Table 1) was comparable (Figure 3A and B).

**Figure 3.**
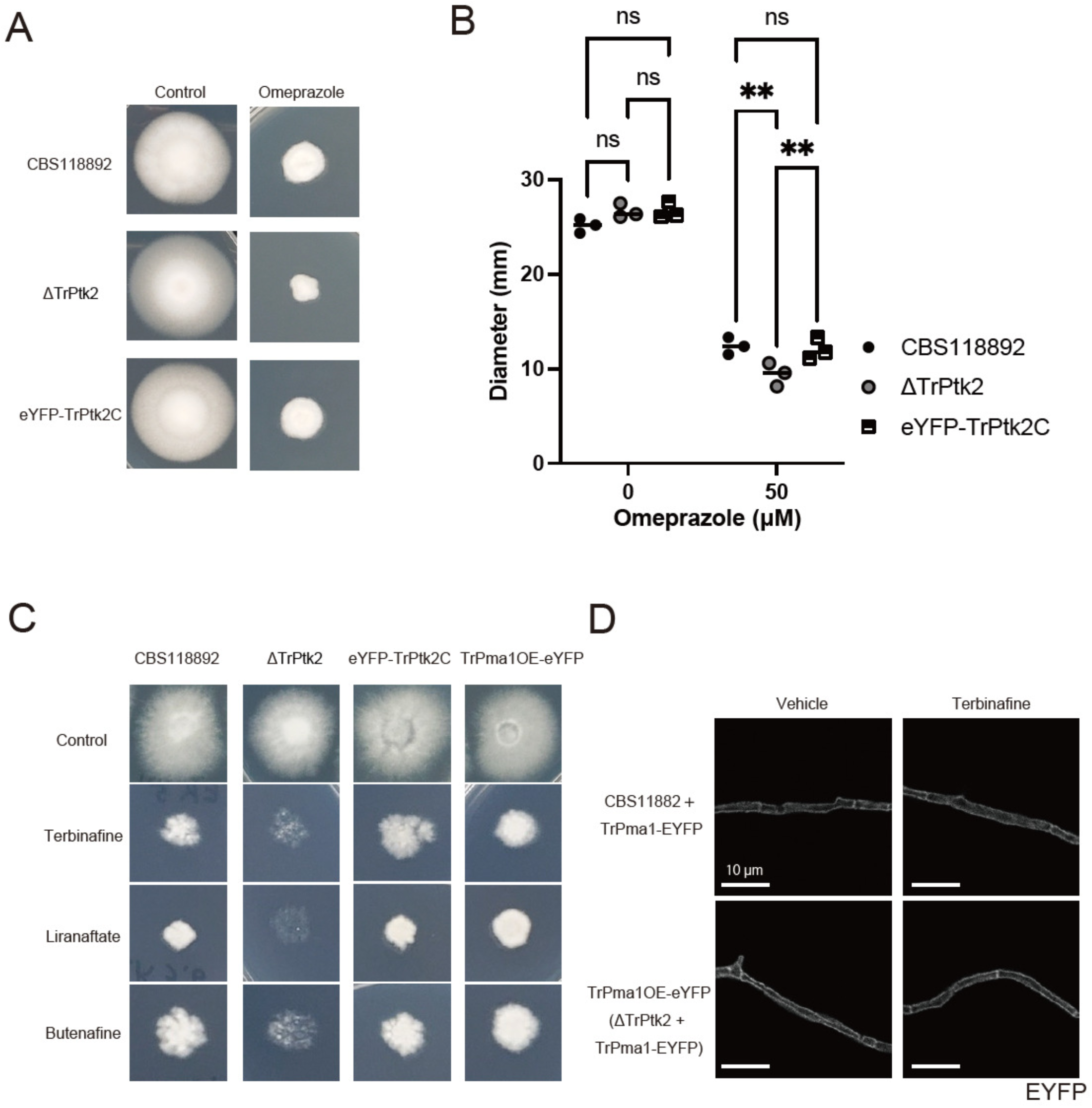
Overexpression of TrPma1 suppresses terbinafine sensitivity of TERG_07844 deletion mutant. (A–B) Spores of CBS118892, ΔTrPtk2, and YFP-TrPtk2C were inoculated on SDA with 0 or 50 μg/mL of omeprazole and incubated for 10 days (A). The diameter of the mycelium was measured after 10 days (B). The dots on the graph represent the diameter of individual samples (n = 3). (C) Spores of strains were inoculated on RPMI 1640 with 5 ng/mL terbinafine, 6.4 ng/mL liranaftate, or 10 ng/mL butenafine and incubated for 14 days. (D) Spores of CBS11882 + TrPma1-eYFP and TrPma1OE-eYFP(ΔTrPtk2 + TrPma1-eYFP) were inoculated on RPMI 1640 and incubated for 2 days or on RPMI 1640 with 0 or 1 μg/mL terbinafine for 3 h. The eYFP signals of the sample were observed under confocal microscopy and shown in white.

If TrPtk2 and TrPma1 function independently, then the genetic inactivation of TrPtk2 in ΔTrPtk2 and the pharmacological inhibition of Pma1 by omeprazole would be expected to have additive effects on mycelial growth. Consequently, there should be no difference in mycelial growth between CBS118892 and eYFP-TrPtk2C on omeprazole-containing medium, just as there is no difference on omeprazole-free medium. However, the diameter of *T. rubrum* ΔTrPtk2 colonies on omeprazole-containing agar was significantly smaller than that of CBS118892 and that of eYFP-TrPtk2C (Figure 3A and B). These results suggest that Ptk2 and Pma1 are part of the same pathway in mycelial growth in *T. rubrum*.

To investigate if TrPma1 functions downstream of TrPtk2, we overexpressed TrPma1 tagged with eYFP at its C-terminus (TrPma1-eYFP) in ΔTrPtk2 and examined whether TrPma1 could complement the terbinafine sensitivity of ΔTrPtk2. On terbinafine-free agar medium (control), *T. rubrum* CBS118892 (parent), ΔTrPtk2, eYFP-TrPtk2C (revertant), and TrPma1OE-eYFP (ΔTrPtk2 overexpressing TrPma1-eYFP) (Table 1) showed similar growth rates (Figure 3C). Mycelial growth was inhibited in *T. rubrum* ΔTrPtk2 not only on agar media containing terbinafine, but also on agar media containing other squalene epoxidase inhibitors, namely liranaftate and butenafine. Conversely, the inhibition of mycelial growth by squalene epoxidase inhibitors was restored in the revertant strain eYFP-TrPtk2C and TrPma1OE-eYFP (Figure 3C). These results suggest that TrPma1 acts downstream of TrPtk2 in the promotion of squalene epoxidase inhibitor resistance.

Since Pma1 functions as a proton pump on the plasma membrane in *S. cerevisiae* (24), TrPtk2 could potentially enhance resistance to terbinafine by regulating the subcellular localization of TrPma1. We overexpressed TrPma1-eYFP in CBS118892 and ΔTrPtk2, then cultivated these strains with or without terbinafine, and examined the subcellular localization of TrPma1-eYFP (Figure 3D). In CBS118892, TrPma1-eYFP localized to the fungal cell surface, as reported for other fungal Pma1 (Figure 3D). The membrane localization of TrPma1-eYFP was not affected in this strain cultured on terbinafine-containing agar medium, indicating that terbinafine does not disrupt the subcellular localization of TrPma1-eYFP. The localization of TrPma1-eYFP on the fungal cell surface was not disrupted in ΔTrPtk2 in the presence or absence of terbinafine. These results suggest that TrPtk2 is not involved in the regulation of TrPma1 subcellular localization and that TrPtk2 promotes terbinafine tolerance by a mechanism other than the regulation of TrPma1 subcellular localization.

### Omeprazole enhances the antifungal activity of terbinafine in both terbinafine- susceptible and -resistant strains

TrPtk2 inhibitors may be effective compounds for combination therapy for dermatophytosis, as ΔTrPtk2 displayed greater sensitivity to terbinafine compared to the terbinafine-susceptible strain CBS118892 (Fig. 1F and G). However, no fungal Ptk2 inhibitors have been identified to date. We hypothesized that pharmacological inhibition of Pma1 might improve dermatophyte sensitivity to terbinafine, since TrPma1 functions downstream of TrPtk2 (Fig. 3C). We assessed growth characteristics of CBS118892 on agar medium containing terbinafine and the fungal Pma1 inhibitor omeprazole. Terbinafine alone had a significant inhibitory effect on the mycelial growth of CBS118892 dermatophytes (Figure 4A and B). Furthermore, the combination of omeprazole and terbinafine resulted in greater inhibition of mycelial growth than either omeprazole or terbinafine treatment alone (Figure 4A and B). These results suggest that omeprazole increases the terbinafine sensitivity of terbinafine-susceptible dermatophyte strains.

**Figure 4.**
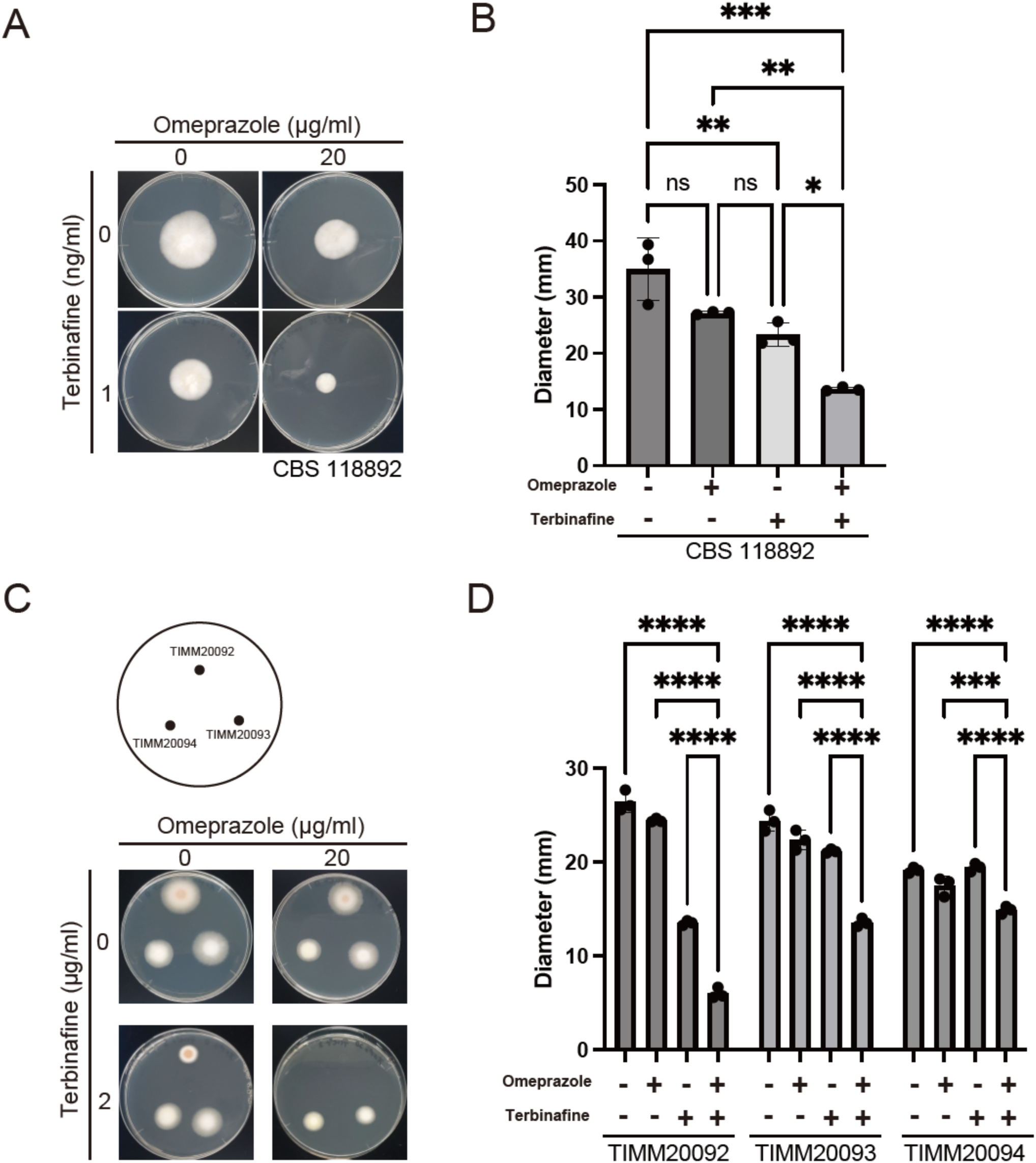
The proton pump inhibitor, omeprazole, enhances the antifungal activity of terbinafine in terbinafine-resistant isolates. (A-B) Combination of omeprazole and terbinafine resulted in greater inhibition of mycelial growth than either omeprazole or terbinafine treatment alone. (A) Spores of CBS118892 were inoculated on SDA with 0 or 20 μg/mL omeprazole and/or 1 ng/mL terbinafine and incubated for 15 days. (B) The diameter of the mycelium on SDA with 0 or 20 μg/mL omeprazole and/or 1 ng/mL terbinafine were measured after 15 days of incubation. The dots on the graph represent the diameter of individual samples (n = 3). (C-D) Decreased terbinafine resistance in terbinafine-resistant isolates in the presence of omeprazole. (C) Spores of TIMM20092, TIMM20093, and TIMM20094 were inoculated on SDA with 0 or 20 μg/mL omeprazole and/or 2 μg/mL terbinafine and incubated for 10 days. (D) The diameter of the mycelium on SDA with 0 or 20 μg/mL omeprazole and/or 2 μg/mL terbinafine was measured after 10 days of incubation. Since TIMM20092 cultured on SDA agar with 20 μg/mL omeprazole and 2 μg/mL terbinafine showed almost no mycelial growth, the diameter was designated as the diameter of the original spot. The dots on the graph represent the diameter of individual samples (n = 3).

The resistance of *T. rubrum* to terbinafine is mainly due to specific mutations in the squalene epoxidase. The mutations L393F and F397L showed the highest minimum inhibitory concentrations (7). We investigated if omeprazole could enhance terbinafine sensitivity in resistant strains. For this purpose, we used clinical isolates of terbinafine-resistant strains that had specific mutations in the squalene epoxidase gene. These included strain TIMM20092 with the F397L mutation, and strains TIMM20093 and TIMM20094, both with the L393F mutation (6). In these terbinafine-resistant strains, both omeprazole and terbinafine exhibited inhibition of mycelial growth individually, except for terbinafine-treated TIMM20094, whose mycelial diameter was comparable with that of the vehicle control (Figure 4C and D). Interestingly, co-administration of terbinafine and omeprazole resulted in more pronounced inhibitory effects than either medicine alone (Figure 4C and D). These results suggest that omeprazole enhances terbinafine sensitivity even in terbinafine-resistant dermatophytes.

## Discussion

The present study suggests that the fungal Ptk2-Pma1 pathway promotes tolerance to squalene epoxidase inhibitors, including terbinafine (Figure 5). Although terbinafine has potent antifungal activity on its own, the finding in this study that inhibition of the TrPtk2- TrPma1 pathway enhances the efficacy of terbinafine is clinically important in terms of overcoming terbinafine-resistant strains. Inhibition of the ATPases, including kinases and proton pumps, has recently emerged as a novel therapeutic approach against drug-resistant dermatophytes (25). Our finding that the antifungal efficacy of terbinafine against terbinafine-resistant dermatophytes is enhanced by omeprazole underscores the importance of this strategy. The veterinary antiparasitic milbemycin has been reported to promote the activity of the antifungal drugs itraconazole and voriconazole via inhibition of the dermatophyte efflux pump MDR3 (26). Omeprazole, which was found to enhance the antifungal effect of terbinafine in this study, has an advantage over milbemycin in terms of clinical applicability as it is a drug approved for human use. This study also suggests that compounds that potentiate the promoting antifungal activity of antifungals can be found by repurposing non-antifungal drugs.

**Figure 5.**
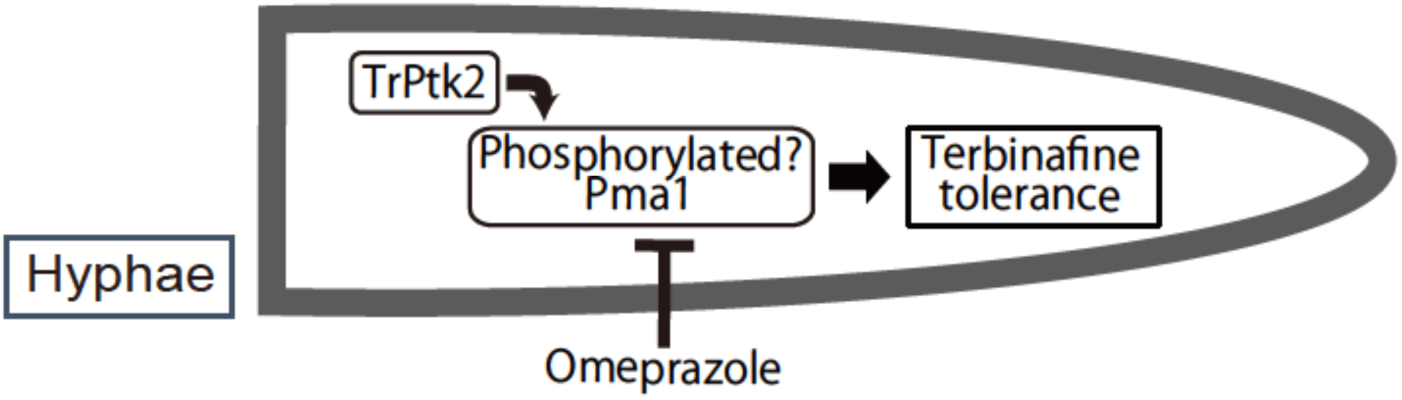
Model of terbinafine tolerance mechanism in *T. rubrum*. Terbinafine tolerance in *Trichophyton rubrum* might be promoted by the phosphorylation of TrPma1 by TrPtk2. The tolerance of *T. rubrum* to terbinafine is decreased by TrPtk2 knockout and omeprazole treatment.

Although our results suggest that TrPma1 functions downstream of TrPtk2 in the promotion of tolerance to terbinafine, TrPtk2 control of TrPma1 was not through regulation of the subcellular localization of TrPma1. The molecular mechanism by which the fungal Ptk2- Pma1 pathway contributes to the terbinafine tolerance remains unknown. In *S. cerevisiae*, Pma1 is phosphorylated by Ptk2 and exports protons from the cell (21, 22). The drug:H^+^ antiporter major facilitator superfamily (MFS) in budding yeast requires the proton gradient that crosses the plasma membrane for drug efflux (27). TrPtk2 may phosphorylate TrPma1, thereby facilitating the formation of the proton gradient necessary for the drug efflux pump to export terbinafine. An efflux pump MDR2 has been identified as a transporter for terbinafine excretion in dermatophytes (28). Since TrPtk2 has also been reported to promote polyamine, Na^+^, and Li^+^ uptake (29–31), it is possible that the TrPtk2-TrPma1 pathway contributes to the acquisition of terbinafine resistance by other mechanisms. Further functional analysis of the TrPtk2-TrPma1 pathway is necessary to better understand terbinafine resistance in dermatophytes. The increased sensitivity to terbinafine of the ΔScPtk2 strain of *S. cerevisiae* lacking Ptk2 demonstrated in this study will also allow *S. cerevisiae* to be used for further studies as a genetic analysis tool alongside studies in *T. rubrum*.

The terbinafine-resistant dermatophyte isolates used in this study have the L393F and F397L substitution mutations in squalene epoxidase (6). The ability of omeprazole to enhance the antifungal activity of terbinafine against these clinical isolates with the major known resistance mutations is critical for therapeutic applications. Furthermore, we found that all three squalene epoxidase inhibitors used in the present study exhibited enhanced antimicrobial activity against ΔTrPtk2 compared to the terbinafine-susceptible parent strain CBS118892. The potential enhanced antifungal activities of squalene epoxidase inhibitors other than terbinafine are important for the further analysis of the function of TrPtk2 as a new medication target and its clinical translation.

## Materials and Methods

### Fungal and bacterial strains and culture conditions

*Agrobacterium tumefaciens* EAT105 (32) was cultured at 28°C in *Agrobacterium* induction medium supplemented with 0.2 mM acetosyringone. Fungal strains used in this study are listed in Table 1. *Trichophyton rubrum* CBS118892, a clinical isolated strain from a patient nail sample was used (12). Terbinafine-resistant *T. rubrum* isolates (TIMM20092, TIMM20093, and TIMM20094) (6) were cultured at 28°C on Sabouraud dextrose agar (SDA; 1% Bacto peptone, 4% glucose, 1.5% agar, pH unadjusted) or 0.165 M MOPS buffered RPMI 1640 agar. *Saccharomyces cerevisiae* BY4741 and YJR059W were purchased from Horizon Discovery Ltd. (California). Parent (BY4741/pYES2-HTH) and ΔScPtk2 (YJR059W/pYES2- HTH) were cultured at 30°C on YPD or synthetic defined medium. Conidia of *T. rubrum* were prepared as described previously (33). pYES2-HTH was purchased from addgene (Massachusetts). For the spot assay, overnight-cultured yeast suspension was diluted with a synthetic defined medium to an optical density of 0.2 (=1.6 × 10^6^ CFU/mL). The suspension was serially diluted, and 3 μL of each suspension was plated onto a synthetic defined agar plate. The samples were incubated at 28°C for 3 days.

### Accession numbers

For comparison of the gene expression of TERG_07844 and other kinase coding genes in *T. rubrum*, we used transcriptome data from the NCBI database (Accession numbers: GSE134406, GSE102872, and GSE110073).

### Plasmid construction

A TERG_07844 (TrPtk2)-targeting vector, pAg1-ΔTrPtk2, was constructed using the following procedure. First, approximately 1.6-kb fragments of the 5’ and 3’-UTR regions of the TERG_07844 open reading frame (ORF) were amplified from *T. rubrum* genomic DNA via PCR with specific primer pairs (F: 5 ′ - GAAGGAGTCTTCTCCTGATCTTCAGCCAAGCAGGG-3 ′ and R: 5 ′ - TCAATATCATCTTCTATCGTCGAGTGGCTTGAGTG-3′; F2: 5′-CGTCATGAATCATCTGCAACTGATCACTGACTGCG−3′ and R2: 5′-GTGAATTCGAGCTCGCCCGCGAAGATCACAGATCA−3′).

The plasmid backbone of pAg and the antibiotic resistance gene cassette were obtained by PCR amplification from the pAg1-3’-UTR of ARB_02021. Finally, these four fragments were fused together using the In-Fusion system (Takara Bio, Inc., Japan).

To construct a vector for TrPtk2 complementation (pCS2-hph-eYFP-TrPtk2), the following steps were performed. First, the antibiotic resistance gene cassette (*hph*) was inserted between the NotI and KpnI sites of pCS2+ N-terminal eYFP, which was generated by inserting the eYFP gene into the BamHI site of pCS2+ (kind gift from S. Kurisu at Tokushima University). Subsequently, the *tef1* promoter (Ptef1) was amplified from the *T. rubrum* genome by PCR with a specific primer pair (F: 5′- GCGGTCGACCCACTAAGACTCCTTCAAGCTCC-3′ and R: 5′-GCGAAGCTTGGTGACGGTGTATTTTTGTGTGG-3′) and inserted between the SalI and HindIII sites of the pCS2+ N-terminal eYFP-derived vector. Finally, the TrPtk2 gene was amplified from *T. rubrum* cDNA via PCR using a specific primer pair (F: 5′- GCTGTACAAGGGATCCATGGCCGGTTCGTCTACAT-3′ and R: 5′-GTTCTAGAGGCTCGATTAGTTGTAGCCATCGCCCA-3′), and the fragment was inserted between the BamHI and XhoI sites of the above vector.

To construct a vector for TrPma1-eYFP overexpression (pCS2-hph-TrPma1-eYFP), the following steps were performed. First, the antibiotic resistance gene cassette (*hph*) was inserted between the NotI and KpnI sites of pCS2+ C-terminal eYFP (kind gift from S. Kurisu at Tokushima University)(34, 35). Subsequently, the *tef1* promoter (Ptef1) was amplified from the *T. rubrum* genome by PCR with a specific primer pair (F: 5′- GCGGTCGACCCACTAAGACTCCTTCAAGCTCC-3′ and R: 5′-GCGAAGCTTGGTGACGGTGTATTTTTGTGTGG-3′) and inserted between the SalI and HindIII sites of the pCS2+ C-terminal eYFP-derived vector. Finally, the Tr*pma1* gene was amplified from *T. rubrum* cDNA by PCR using a specific primer pair (F: 5′- TCTTTTTGCAGGATCGCCACCATGGCCGACCACGCAGCC-3′ and R: 5′-CCTCTAGAGGCTCGAGGTGCGCTCTTCTCGTGCTG-3′), and the fragment was inserted between the BamHI and XhoI sites of the above vector.

### Transformation of *T. rubrum*

The *A. tumefaciens*-mediated transformation (ATMT) technique was used to alter *T. rubrum*, as previously described (13–15). PCR was used to assess the intended transformants and pure genomic DNA. A Quick-DNA Fungal/Bacterial Miniprep Kit (Zymo Research, California) was used to extract the total DNA. The T-01 system (TAITEC, Japan) with 5-mm stainless steel beads was used to perform a study on the collision of beads with fungal cells. For confirmation, PCR was conducted using two primer pairs (Primer 1: 5′- GCTTCTCCATCCCTGCTGTT-3′, Primer 2: 5′ATTCGTCTGCAAGGGGACAG-3′, Primer 3: 5′- AGAAGATGATATTGAAGGAGCACTTTTTGGGCTT-3′, Primer 4: 5′- AGATGATTCATGACGTATATTCACCG-3′)

### Fluorescent microscopy observation

CBS118892 + TrPma1-eYFP or ΔTERG_07844 + TrPma1-eYFP strains were seeded with 1–5 × 10^6^ spores on sterile cover glasses and placed in a 12-well plate. They were then incubated with 500 μL of SD liquid medium at 28°C overnight. On the second day, the SD medium was replaced with fresh medium, and the spores were further incubated at 28°C overnight. On the third day, the supernatant was removed, and the cells were cultured in RPMI 1640 with or without terbinafine for 3 h. The sample was fixed with 4% paraformaldehyde (PFA, Nacalai Tesque, Japan) at room temperature for 15 minutes. The samples were washed three times with PBS and mounted on glass slides using Aqua-Poly/Mount (Polysciences, UK). The stained cells were observed using a confocal microscope system (AX, Nikon, Japan).

### Phylogenetic tree analysis

The evolutionary history was inferred using the maximum likelihood method and the Whelan and Goldman + Freq. model. The tree with the highest log likelihood (−13606.41) was used. The percentage of trees in which the associated taxa clustered together was shown below the branches. The initial tree(s) for the heuristic search were obtained automatically by applying the Neighbor-Join and BioNJ algorithms to a matrix of pairwise distances that were estimated using the JTT model, and then selecting the topology with the superior log likelihood value. A discrete Gamma distribution was used to model the evolutionary rate differences among the sites (5 categories [+G, parameter = 5.3335]). The tree was drawn to scale, with branch lengths measured in the number of substitutions per site. This analysis involved eight amino acid sequences. There was a total of 1,163 positions in the final dataset. The evolutionary analyses were conducted in MEGA11 software.

### Statistical analysis

The means of the two groups were compared using Student’s *t*-test. For three or more groups with a single variable, one-way analysis of variance (ANOVA) with Tukey’s post hoc test was conducted. For means of three or more groups with two variables, two-way ANOVA with Tukey’s post hoc test was performed. Prism 9 software (GraphPad Software, Boston) was utilized for these statistical analyses. Statistical significance was defined at a *p* value of <0.05.

## Abbreviations

SD: Sabouraud dextrose
SDA: Sabouraud dextrose agar

## Acknowledgments

The authors thank Dr. S. Kurisu at Tokushima University for providing the pCS2+ N-terminal and C-terminal eYFP plasmids, K. Maru, H. Shimokuri, S. Yahagi, H. Uga, and N. Hori for their technical help. This work was supported by the Japan Society for the Promotion of Science and Takeda Science Foundation.

